# Overexpression of Zinc finger (GpZF) promotes drought tolerance in grass pea (*Lathyrus sativus*)

**DOI:** 10.1101/2021.10.19.464954

**Authors:** Mitra Parsa, Mona Kashanchi, Amineh Zeinali, Elaheh pourfakhraei

## Abstract

The genes encoding Cys2/His2-type zinc finger proteins constitute a large family in higher plants consisting of a family of plant transcription factors. GpZF encodes a Cys2/His2-type zinc finger protein. The purposes of this study are to elucidate further the functions of a novel zinc finger transcription factor in the grass pea (GpZF) gene involved in the drought stress response in the grass pea (*Lathyrus sativus*) and to investigate its biochemical and physiological parameters under stress conditions. GpZF was expressed in grass pea. Relative gene expression analysis showed the GpZF gene in independent transgenic lines was more under drought mild and severe treatments (50% and 25% field capacity-FC). Furthermore, overexpression of this gene in grass pea results in more relative water content, free proline, and soluble sugars than the wild-type (WT) plants under 50% and 25% FC stresses. Moreover, in 25% FC, the independent transgenic lines revealed an increase in survival rates and dry weight than the WT plants. GpZF thus implies the positive role in drought stress tolerance in *Lathyrus sativus*. In conclusion, the transgenic grass pea plants generated in this study could be used to farm arid areas.

**Highlight:** GpZF is a positive regulator in drought stress tolerance in grass pea.

## 1. Introduction

Environmental stresses, such as drought, high salinity, and low temperature, alter plant water status and severely limit plant growth and productivity primarily thanks to the photosynthetic shortage, osmotic stress restraints on plant processes, and nutrient availability interference (Chen *et al*., 2010; Dixit *et al*., 2016; Hu *et al*., 2009; Ni *et al*., 2009; Varshney *et al*., 2009).

Adaptation to these stresses in plants occurs at molecular, cellular, physiological, and biochemical levels. At the molecular level, transcription factors (TFs), as regulatory genes play critical roles in responding to abiotic stresses (Wang *et al*., 2014). One of the most prominent transcription factor gene families is the Zinc-finger (ZF) family (Hall, 2005). This group of TFs participate in many metabolic pathways, various biological functions, such as plant growth and development (Chrispeels *et al*., 2000; Kubo *et al*., 1998; Sakai *et al*., 1995), phytohormone (Molnár *et al*., 2002) and biotic and abiotic stress responses (Ciftci-Yilmaz & Mittler 2008; Lippuner *et al*., 1996; Mukoko Bopopi *et al*., 2010; Takatsuji 1998; Tian *et al*., 2010; van der Krol *et al*., 1999). The Cys2/His2-type zinc-finger is identified as DNA-binding motifs in eukaryotic transcription factors TFIIIA-type finger, which typically have a QALGGH motif within the zinc finger protein (ZFP) domain. However, it has been recognized several plants C2H2-type ZFPs without the QALGGH motif.

Grass pea (*Lathyrus sativus* L.) is an annual and diploid (2n = 2x = 14) pulse crop belonging to the tribe vicieae in the family Fabaceae (Yang *et al*., 2014). Despite its tolerance to drought, grass pea is not affected by excessive precipitation and flooding, including poor soils and heavy clays (Almeida *et al*., 2015; Jiang *et al*., 2013; Urga *et al*., 2005; Yang *et al*., 2014). It is an excellent candidate crop for human diets and animal feeds in drought-prone areas because of promising starch and protein (Jiang *et al*., 2013; Yang *et al*., 2014). Therefore, breeding drought-tolerant grass pea cultivars is imperative to adapt to environmental stresses.

It has been reported that TFs play a critical role in the regulation of many stress-responsive genes. As well, they can change the downstream regulation of gene expression and signal transduction in stress-response pathways.

ZFP36, has been shown to be induced by drought, oxidative stress, and regulated for the cross-talk between NADPH oxidase, H2O2, and MAPK in ABA signaling which lead to tolerant in rice plants (Zhang *et al*., 2014). ZFP185 has been displayed to be regulated plant growth and stress responses by affecting GA and ABA biosynthesis in rice (Zhang *et al*., 2016). ZAT18, a nuclear C2H2 zinc finger protein, functions as a positive regulator in the plant response to drought stress in Arabidopsis (Yin *et al*., 2017).

They then are good candidates for the breeding stress-tolerant crop. Because zinc finger is a major transcription factor that responds to abiotic stresses (salt, dehydration, and cold), in the present study, we report on the identification of grass pea of zinc finger (GpZF), a zinc finger TF in grass pea.

We found that the zinc finger Overexpressing showed to enhance drought resistance in transgenic grass pea. Moreover, biochemical and physiological parameters relating to stress were evaluated to demonstrate the function of the zinc finger.

## 2- Materials and Methods

### 2-1- Plant material, growth conditions and stress treatments

Healthy and mature seeds of grass pea (*Lathyrus sativus*), provided by the Research Institute of Forests and Rangeland was used as material for subsequent procedures.

After sterilizing the seeds, they were placed for germination on MS medium. The seedlings were grown at 26±1°□ under continuous light conditions for 5 to 7 days until the axillary buds are prominent. The axillary meristem explants were subcultured and used for transformation purposes.

The transgenic lines and wild-type plants were propagated from the cotyledon node. The transgenic plants were taken from the tubes carefully, washed and dipped in a 0.5% thiram solution and transferred to 8-cm-diameter pots containing 2 to 4 mm of sand followed by 20-cm-diameter pots containing the potting mixture. Plants were transferred to a greenhouse with 24/18° □day/night temperatures and allowed to grow until maturity. The wild-type plants’ process was similar to the transgenic ones except for the 0.5% thiram solution step.

For drought treatment, soil-grown 12 day-old plants were subjected to progressive drought by two different water treatments (50 and 25% of field capacity – FC) was applied based on FC. The FC was determined by the gravimetric method following Souza *et al*. (2000) methodology. The drought-stressed plants compare with well-watered plants as a control. After harvesting drought-stressed and unstressed plants at the same time of the day, these samples were immediately frozen in liquid nitrogen and stored at −80°C before RNA isolation.

### 2-2- Phylogenetic Analysis

Multiple sequence alignment was performed using the National Center for Biotechnology Information (NCBI) constraint-based multiple alignment tools (https://www.ncbi.nlm.nih.gov/tools/cobalt/cobalt.cgi). A phylogenetic tree was constructed with the aligned plant C2H2 zinc finger proteins using MEGA version 5.0 via the neighbour-joining method.

### 2-3- Transgenic Plasmid Construction

The full-length cDNA of *GpZF* was obtained from *Lathyrus sativus* using gene-specific primers: the forward primer 5‘TCTAGAATGCCAACAGTGTGGTTTTC3’ and reverse primer 5‘TCCAGA TCATGGTTTGCAAATTACGA3’. To generate *GpZF*-overexpressing of transgenic grass pea, the *GpZF* complementary DNA (cDNA) was subcloned into the grass pea transformation vector, pBI121 under the control of the cauliflower mosaic virus (CaMV) 35S promoter. The recombinant vector was transformed into Agrobacterium strain C58. Grass pea (*Lathyrus sativus*) was transformed using the cotyledon node method (Wang, 2006).

### 2-4- RNA purification and cDNA synthesis

According to the manufacturer’s protocol, total RNA from each sample was extracted using the Ribospin Plant kit (Gene All, Korea). The samples’ concentration and integrity were examined by nanodrop (Thermo Fisher Scientific, 2000, USA) and 1.2% agarose gel electrophoresis. The first-strand cDNA was synthesized based on 1µg total RNA of each sample using the M-MLV reverse transcription system (Ferementas, USA) following the method provided by the manufacturer’s instructions.

### 2-5- Validation of candidate genes using qPCR analysis and efficiency

Leaves of WT plants and independent transgenic lines (11, 22 and 50) exposed to drought treatments (50% and 25% field capacity) were used qPCR analysis. Real-time quantitative PCR was conducted with Real-time PCR System (Qiagen, Germany), using Syber Green qPCR Master Mix2x (Ampliqon, Denmark) to monitor dsDNA synthesis. An endogenous ß tubulin was used as an internal standard for the drought stress marker gene. Primers were designed by NT Vector software (table 1). Relative expression levels were calculated using the delta threshold cycle (Ct) methods and 2^□ Δ ΔCT^ (Livak & Schmittgen, 2001).

**Table 1.**
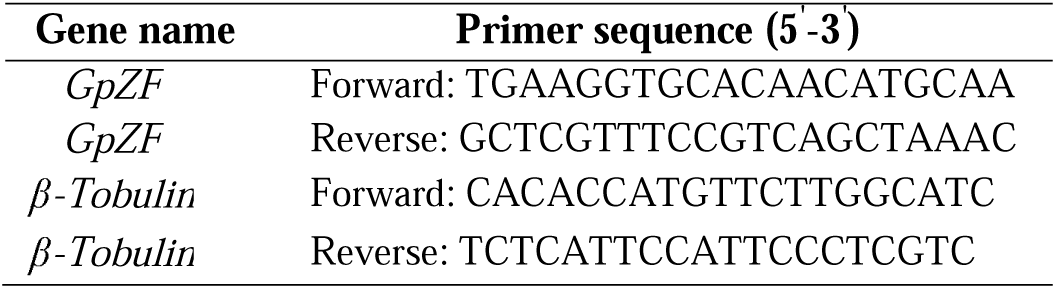
gene-specific primers used for quantitative real-time PCR

### 2-6- Measurement of free proline and total soluble sugar contents

Free proline content of drought-treated WT and transgenic grass pea plants was measured through spectrophotometer according to the method of (Bates *et al*., 1973). The content of proline (µg/g FW) was quantified by the ninhydrin method. The phenol–sulfuric acid method was used to assess plants’ total soluble sugar content (Dubois *et al*., 1956). Finally, the absorbance of proline and sugar content was determined at 520 nm and 485 nm with a spectrophotometer respectively (WPA, Sweden).

### 2-7- Measurement of relative membrane permeability and relative water content

Membrane permeability was measured according to (Bajji *et al*., 2002), with some modification. In this method, three discs in the leaf segments from seedlings were infiltrated and maintained in water for 25 min and 2 h respectively. The electrical conductivities (C1) of the obtained solutions were then determined. Next, boiling the leaf segments in deionized water was occurred for 15 min and the conductivities (C2) of the resulting solutions were measured. The relative electrolyte leakage was calculated and evaluated by the values of C1 to C2 (C1/C2).

The fresh and dry leaves of grass pea plants were weighed. The relative water content (RWC) (Pieczynski *et al*., 2013) was evaluated before and after the drought treatment.

### 2-8- Statistical analysis

The root length, shoot weight, and survival rate were calculated three times repeats. For relative membrane permeability, relative water content, free proline, total soluble sugars, samples were taken in three biological replicates at the indicated time after treatment. To discriminate significant differences, the data were interpreted by analysis of variance (ANOVA), and the t-test was used for determination of the least significant difference (LSD) of means.

## 3- Results

### 3-1- Bioinformatic analysis of the gene GpZF

The GpZF gene, containing a complete ORF of 1274 bp, was cloned from total RNA prepared from *Lathyrus sativus* seedlings using RT-PCR. This gene was homologous to C2H2-type ZFPs of Trifolium (Fig. 1A). These genes contain one Cys -x(2,4)-Cys -x(3)-[LIVMFYWC]-x(8)-His -x(3,5)-His C2H2-type zinc finger, without a QALGGH motif. A phylogenetic tree was constructed by the Maximum Likelihood method with MEGA5.0 to investigate the evolutionary relationship among plant C2H2-type ZFs (Fig. 1B). The results demonstrated that GpZF was clustered with C2H2-type zinc fingers Trifolium. The upstream cis-acting regulatory element of GpZF was analyzed and found that stress- and defence-related elements, such as W-boxes (position-134 to-139), TC-rich repeats (position-1131 to - 1140), and CGTCA (position -855 to -860), were enriched in the promoter region of GpZFp, indicating that GpZF is a stress-related mRNA.

**Fig. 1.**
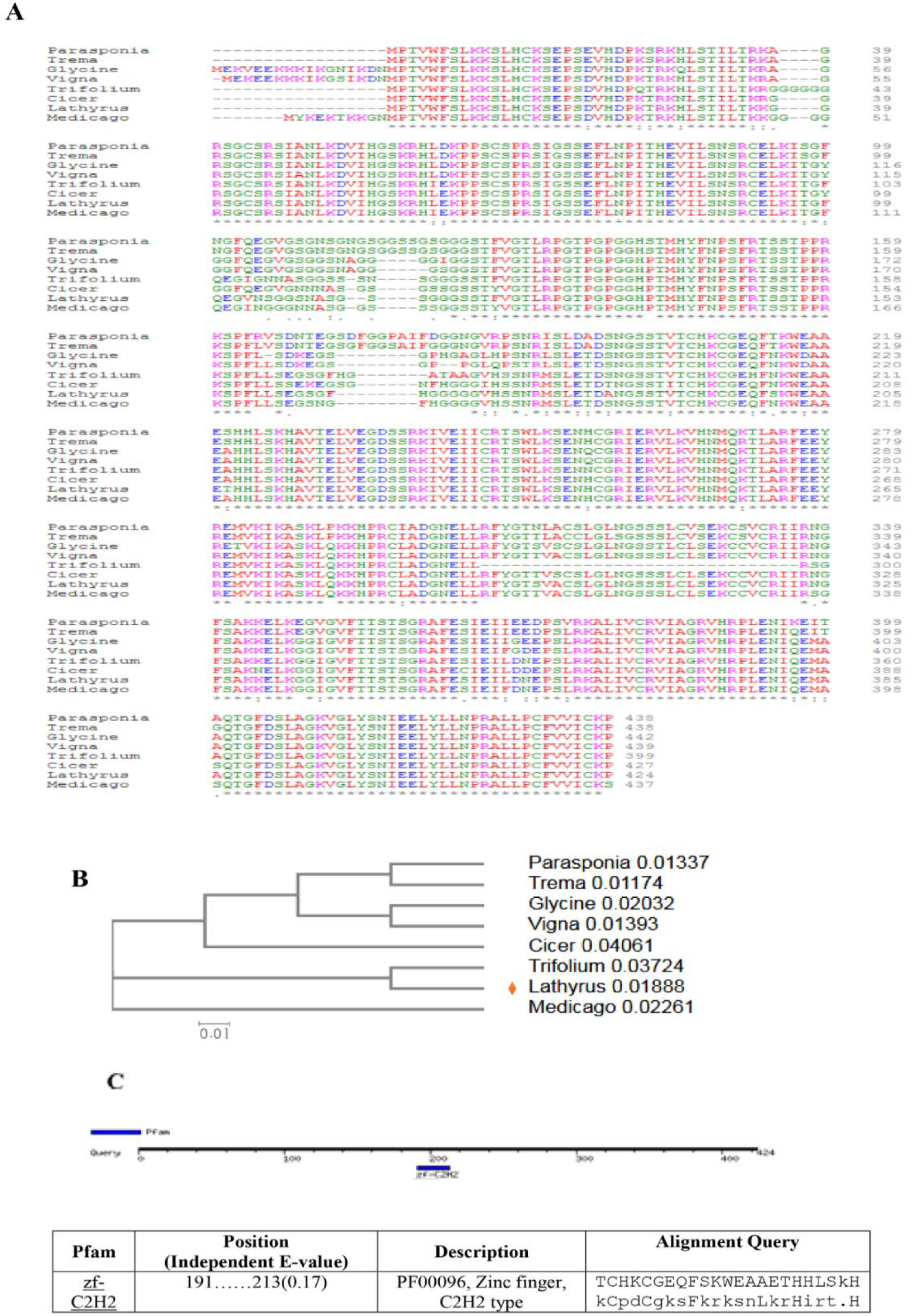
Bioinformatic analysis of GpZFp. A Multiple sequence alignment of amino acid sequences of GpZF using ClustalW. B The phylogenetic tree of plant stress-responsive C2H2-type ZFPs. C The position of C2H2 zf in sequence.

### 3-2- Identification transgenic grass pea plants

To elucidate the function of the GpZF in grass pea, the 35S: GpZF construct (Fig. 2A) was transformed into *Lathyrus sativus* via Agrobacterium. The lines of transformed explants were identified by PCR first, and 52 independent lines were PCR positive (data not shown). The percentage of PCR regenerated lines is 36.6 %. Semi-quantitative RT-PCR was conducted to evaluate the expression level of GpZF in these independent lines. The expression level of GpZF in three different independent transgenic lines; 11, 22 and 50 was significantly higher than that in the wild-type plants (Fig. 2B), which qRT-PCR analysis confirmed the expression level of this gene in mentioned lines. These were chosen for further analysis subsequently (Fig. 2C). Being an annual legume, T0 generation of grass pea seedlings were used to verify drought tolerance of GpZF.

**Fig. 2.**
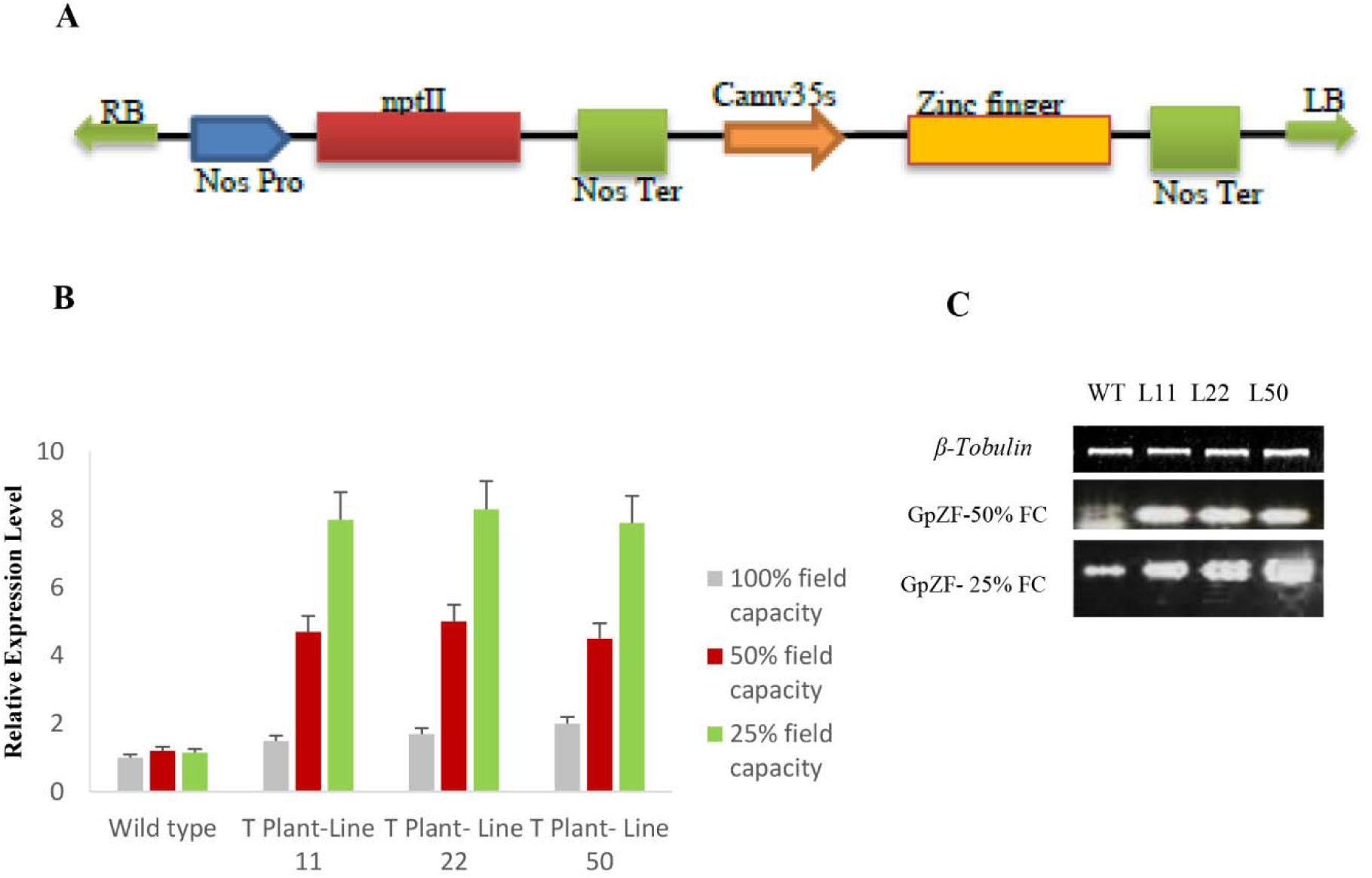
A. Map of the pbI-GpZF plasmid. The GpZF gene was under the control of the CaMV 35S promoter. B and C. Analysis of GpZF transcript level in transgenic grass pea and WT plants by qRT-PCR and real-time RT-PCR. WT: non-transgenic plants, and 2, 11 and 22: independent transgenic lines. B. Real-time PCR analysis of GpZF expression in the presence and in the absence of drought (50% and 25% field capacity); values indicate means of three biological replicates. Significant differences indicate by Student’s t-test and P < 0.05.

### 3-3- Overexpression of GpZF on growth conditions in transgenic grass pea

After regeneration of grass pea (Fig. 3A, B, C and D), the T0 generation of the putative transformants for the incorporated gene was tested.

**Fig. 3.**
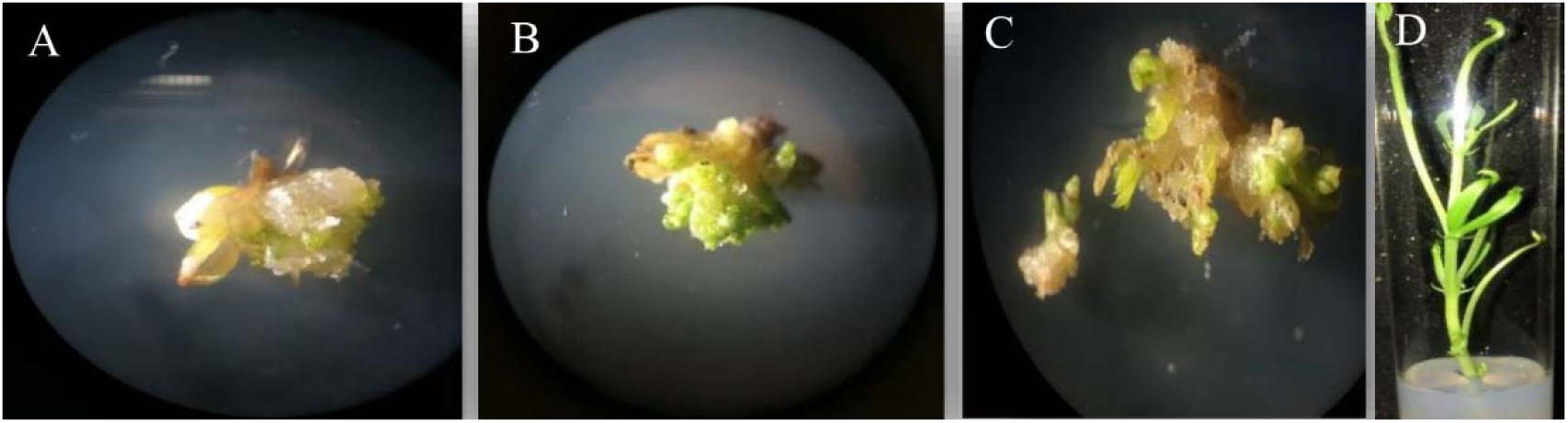
Regeneration of adventitious shoot buds and plants from axillary meristem explants of transgenic grass pea. A Callus induction from axillary explant B and C Somatic Embryogenesis D Elongation of shoot.

Under normal growth conditions, overexpression of GpZF does not lead to any significant observable effects in plant growth habits. However, after drought treatment (50% and 75% field capacity), leaves of wild-type plants showed severe wilting and chlorosis, whereas transgenic lines grew well with leaves slightly yellow. In drought stress, transgenic lines grew better than wild-type plants. The shoot height of transgenic lines was considerably higher than that of wild-type plants (Fig. 4A, B). Furthermore, the survival rate of transgenic lines was significantly higher than that of wild-type plants (Fig. 4C).

**Fig. 4.**
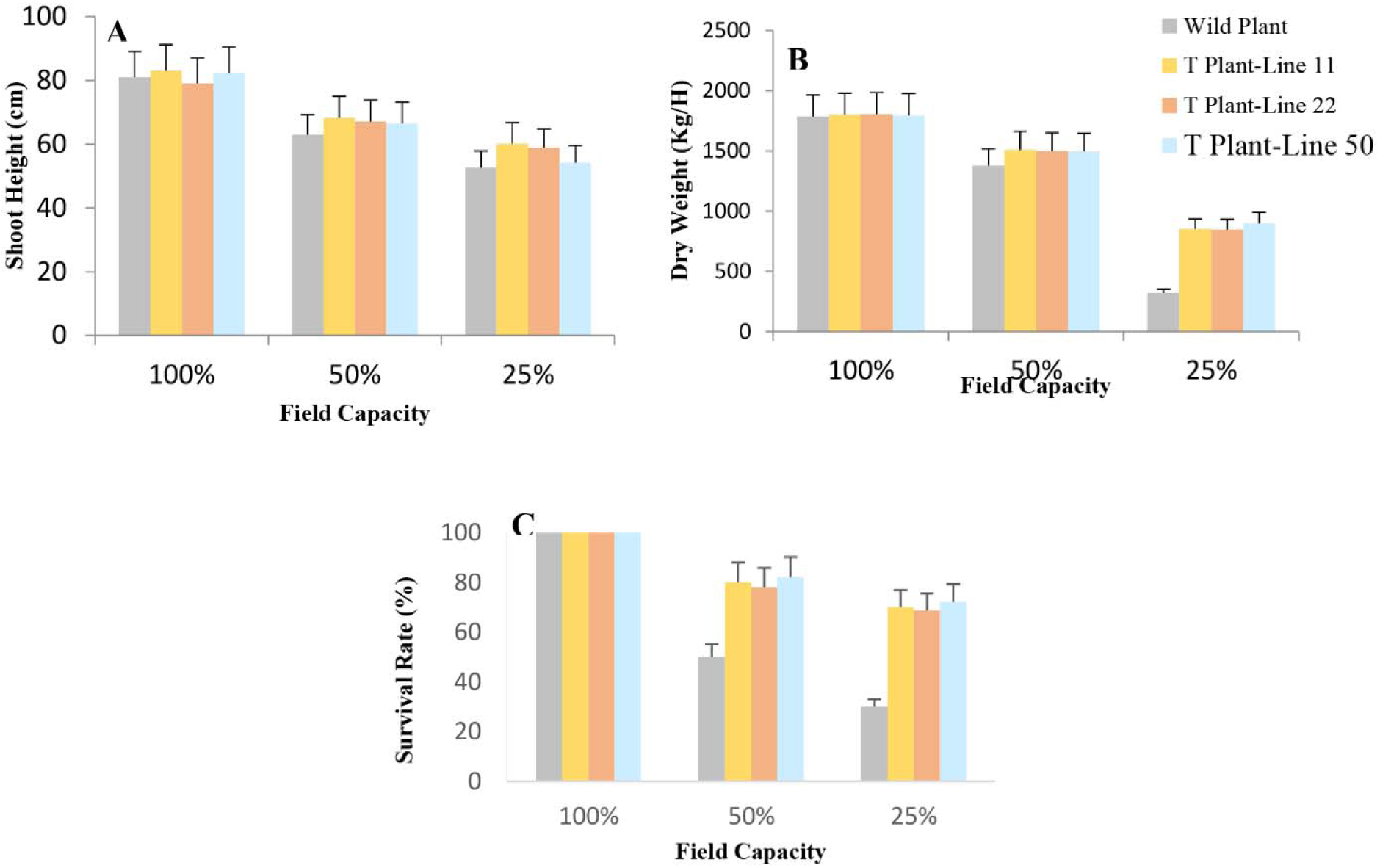
Drought stress tolerance of GpZF in transgenic grass pea. A Shoot height of plants under control conditions and drought treatments. B. Dry weight of plants in the presence and in the absence of drought. C. Survival rate under drought stress

Under drought conditions, the relative water content and the relative membrane permeability of wild-type plants was more remarkable than those of the lines overexpressing GpZF (Fig. 5A and B).

**Fig. 5.**
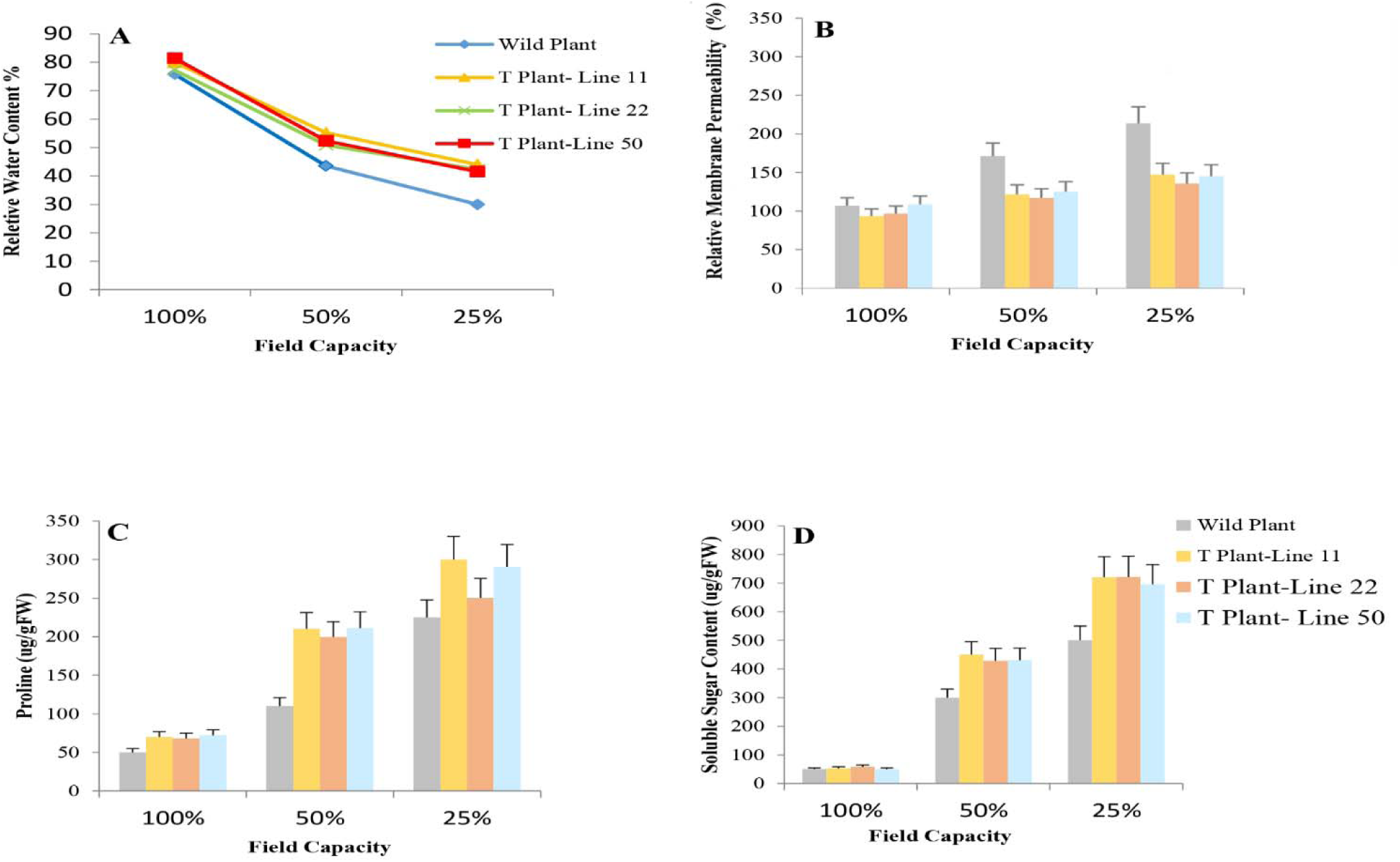
Drought stress tolerance in wild-type and transgenic grass pea A. Relative water content, B. Relative membrane permeability, C. Free proline, D. Soluble sugar content.

As a result of abiotic stresses, plants can cumulate more osmolytes compatibility, such as free proline and soluble sugars as osmoprotectants (Irigoyen *et al*. 1992). Therefore, proline and soluble sugar contents were analyzed in transgenic and wild-type plants. There was no significant difference between wild-type and transgenic lines under the control conditions; however, after drought treatment, compared with the wild-type plants, independent transgenic lines showed markedly higher free proline and soluble sugars (Fig. 5C, D).

## 4. Discussion

Transcription factors are the important components of the complicated gene regulatory networks that can regulate the expression of the amount of various stress-responsive genes to deal with stress (Kiełbowicz-Matuk, 2012; Lan *et al*., 2017). The C2H2-type zinc finger transcription factors were caused to be involved in plant development and have various adaptive responses to abiotic stresses such as drought. In the present study, we characterized a new C2H2 zinc finger transcription factor ZF (GpZF) from grass pea. Sequence analyses revealed that GpZF had an identity with other C2H2 zinc finger proteins without QALGGH amino acid sequences. Lack of conserved QALGGH motif has been previously been shown to enhance Arabidopsis tolerance to cold stress (Luo *et al*., 2014). However, studies on GsZFP1 gene transformation without the typical QALGGH motif in Arabidopsis revealed that it played an important role in withstanding cold and drought stresses (Luo *et al*., 2012), suggesting that the QALGGH motif in the GsZFP1 protein was not imperative for adaptation to abiotic stress in wild soybean. In our study, GpZF also did not have a plant-specific typical QALGGH motif, meaning that this motif was not vital for plant response to abiotic stresses in grass pea.

Although decreased gas exchange results in the reduction of photosynthetic production during stomatal closure, decreased transpiration can reduce water loss from leaves (Ciftci-Yilmaz & Mittler, 2008). Therefore, this process leads to growth inhibition. In drought conditions, ABA alteration of guard cell ion transport, increasing stomatal closure and preventing stomatal opening, which reduces water loss (Kim *et al*., 2011). A sensitivity to ABA is not necessarily coincided with an increase in stress tolerance in plants (Ren *et al*., 2005). Stomata openings also not always occurred with a rising sensitivity to drought and water loss rates. For example, in Arabidopsis, the percentage of open stomata is much less in the WT plants than the LLA23 over-expression gene under stress, while transgenic plants have a declined rate of water loss and exhibit increased drought (Yang *et al*., 2005). In this work, there is meaningful difference in growth and development between the independent transgenic lines and the controls under severe drought stress. It has been demonstrated that lower stomata density and size do not mean biomass accumulation and lower CO2 assimilation (Yoo *et al*., 2009). For example, alteration in stomata development did not affect CO2 assimilation and biomass accumulation (Yoo *et al*., 2009). stomata density and pore aperture operate stomatal conductance (Hetherington & Woodward, 2003; Nilson & Assmann, 2007). It is shown that the average stomata density in the GsZFP1 overexpression lines was similar to that found in WT plants. For most C3 plants, the net CO2 assimilation rate saturates as stomatal conductance enhances as a result of non-stomata limitations, such as the regeneration of ribulose 1,5-bisphosphate (Zeiger & Field, 1982). Therefore, a difference in stomata development cannot have a coincidental effect on CO2 assimilation. Overexpression of PtXERICO as a focal point in woody perennials improved drought tolerance by upregulation of endogenous ABA level (Kim *et al*., 2020).

At the cellular level, membranes are thought to be the site of primary physiological injury (Blum 1988) and an early event in plant response to abiotic stress. Our results showed that under drought, the degree of membrane injury of wild type plants was more than transgenic plants.

Besides, the GpZF could regulate many biological processes to deal with drought, including proline and soluble sugar synthesis. It is proven that the accumulation of soluble sugars is considerably correlated to the drought tolerance in plants (Hoekstra, 2001), and usually, the measurement of proline content is relatively dependent on the sugar levels (Larher *et al*., 1993).

Sun *et al*., revealed that suffering plant from salt stress, ZFP179 might increase the expression of stress defence genes such as OsP5CS and follow by the accumulation of free proline, soluble sugars (Sun *et al*., 2010).

At this work, transgenic grass pea accumulated higher levels of proline and soluble sugars that function as osmolytes than the WT plants under drought stress conditions. Maintaining a high RWC occurred as a result of the increased soluble sugar and proline levels. In comparison with WT plants, the transgenic plants illustrated high RWC content.

## Abbreviations

GpZF: grass pea Zinc Finger
FC: Field capacity
TF: transcription factor
qPCR: quantitative real-time polymerase chain reaction
WT: wild-type
RWC: relative water content

## Acknowledgments

The work was supported and funded by the ACECR and the Research Institute of Applied Science. In addition, the authors thank the Research Institute of Forests and Rangeland and the agricultural organization of Golestan province for providing the grass pea seeds.

## Conflict of interest

The authors declare no conflict of interest.

## Author contributions

Mitra Parsa designed the project and analyzed and interpreted the data; Mona kashanchi and Amineh Zeinali performed the experiments; Elaheh Pourfakhraei helped analyze the data and write the paper.

## Data availability

All data supporting the findings of this study are available within the paper.

## Notes

### Competing Interest Statement

The authors have declared no competing interest.

